# Implications of diabetes mellitus on pathophysiology of Tuberculosis: Role of AGE-RAGE Axis and Glycated Proteins

**DOI:** 10.1101/2024.12.15.628594

**Authors:** Sudhasini Panda, Alankrita Singh, Diravya M Seelan, Alisha Arora, Gunjan Dagar, Abdul S Ethayathulla, Kalpana Luthra, Anant Mohan, Naval K Vikram, Neeraj Kumar Gupta, Mayank Singh, Archana Singh

## Abstract

Chronic hyperglycemia in diabetes mellitus promotes the formation of advanced glycation end products (AGEs) via non-enzymatic glycation of plasma proteins. AGEs and their receptor RAGE are central to diabetic complications, as they activate inflammatory pathways and oxidative stress. We studied the AGE-RAGE axis in uncontrolled diabetic individuals having pulmonary TB (PTB+DM), uncontrolled diabetic individuals with no infection (DM), and pulmonary TB patients (PTB). Levels of AGEs and RAGE were measured across these groups and compared to controls. The results showed significantly higher AGEs and RAGE levels in the diabetic milieu, reflecting chronic hyperglycemia. These AGEs were shown to be associated with impaired macrophage function, evidenced by reduced phagocytic capacity. Increased AGEs correlated with disease severity in tuberculosis patients co-affected by diabetes, indicating compromised immune responses. RAGE expression also showed a higher trend in TB patients, although not statistically significant, potentially influencing infection outcomes. Glycated calmodulin was also elevated in diabetic and TB-diabetes comorbid patients and negatively correlated with nitric oxide (NO) levels, suggesting that calmodulin glycation hinders NO production in diabetic conditions. Molecular dynamic studies revealed that glycosylation disrupts calmodulin’s interaction with inducible nitric oxide synthase (iNOS), potentially explaining the decreased NO levels in TB-diabetes comorbid patients. These findings underscore the detrimental impact of AGEs and RAGE in diabetic immune dysregulation, highlighting the need for therapeutic strategies to mitigate their effects on macrophage function and restore immune responses.

**Highlights:**

- We conducted this study to examine how hyperglycemia-driven AGEs and RAGE contribute to immune dysfunction and disease severity in diabetic and TB-diabetes patients.
- High AGEs and RAGE impairs macrophage function, decreasing phagocytic capacity and worsening disease severity in TB-diabetes comorbidity.
- Glycosylation disrupts calmodulin’s interaction with iNOS, contributing to lower NO levels and weakened immune responses in TB-diabetes comorbidity.
- The AGE-RAGE axis should be studied further for these comorbid conditions to assess and improve macrophage function and restore immune responses.

---

Diabetes is a metabolic disorder, characterized by hyperglycemia and its sequelae. Patients with diabetes mellitus (DM) have infections more often than those without DM. Impact of diabetes on TB control is particularly alarming, as an estimated 11% of all global TB deaths are attributable to diabetes (1). The increased incidence of TB in people with DM appears to be multifactorial (2). Chronic inflammation and oxidative stress due to long standing hyperglycemia may affect TB infection outcome. During long standing hyperglycaemic state in diabetes mellitus, glucose forms covalent adducts with the plasma proteins through a non-enzymatic process known as glycation and form advanced glycation end products (AGEs) which play an important role in the pathogenesis of diabetic complications (3,4). It can modulate immune response as well by binding to its receptor, RAGE and initiation of signalling cascade leading to inflammation, release of pro-inflammatory molecules and free radicals(5,6). The formation of AGEs or RAGE ligands might be relevant to TB-DM comorbidity because RAGE ligands are known to accumulate to a greater level with DM and also in the presence of chronic inflammation, such as that which occurs in TB. It is possible that RAGE ligands may accumulate at a faster rate in people with TB-DM, given the convergence of hyperglycemia and chronic inflammation. RAGE ligand upregulation may alter immune cell function and lead to a prolonged proinflammatory response through RAGE-mediated activation of nuclear factor kappa-light-chain-enhancer of activated B cells (NF-kB) (7,8). Therefore, in the present study, we have tried to assess the effect of AGE-RAGE in uncontrolled diabetic patients having tuberculosis.

Additionally, it is known that glycation of proteins under chronic hyperglycemic condition can lead to structural changes and hence affect their cellular functions. Two such proteins of our interest include calmodulin and inducible nitric oxide synthase (iNOS). iNOS is required for the production of reactive nitrogen species (RNS) which helps in bacterial killing. This enzyme which usually occurs in monomeric form is dimerized by Calmodulin (CAM) which initiates the dimerization of iNOS and results in active form of iNOS and hence the production of RNS(9,10). Therefore, we have also checked for the levels and association of glycated iNOS and calmodulin in the present setting. We performed Molecular Dynamics Simulations of CaM-iNOS-FMN complex and assessed their binding efficiency upon glycosylation.

## METHODS

### Study participants

This cross-sectional study was conducted at AIIMS, New Delhi, with participants divided into four groups: pulmonary TB (PTB), type 2 diabetes (DM), PTB + DM, and healthy controls. The study enrolled 65 newly diagnosed, treatment-naive PTB patients, defined by clinical diagnosis, sputum smear, culture, or Gene Xpert positivity. Exclusion criteria included extra-pulmonary or drug-resistant TB, HIV, other significant diseases, pregnancy, and concurrent illnesses.

The DM group comprised 51 uncontrolled type 2 diabetic patients (HbA1c >7.5%) without other disorders or TB. PTB + DM included 50 newly diagnosed PTB patients with uncontrolled type 2 diabetes (HbA1c >7.5%), meeting criteria similar to PTB and DM groups. The healthy control group included 50 individuals without TB, diabetes, or other disorders. All participants (aged 18–50 years) were sex-matched, with record of their BMI. Ethical approval (IECPG-374/28.09.2017) was obtained, and participants provided informed consent. The demographic information is available in our previous published paper (11).

### Generation of monocyte-derived macrophages (MDMs)

Peripheral blood mononuclear cells (PBMCs) were isolated using Ficoll gradient separation, followed by monocyte isolation via plastic adherence. 1–2 million PBMCs were plated in 12-well plates with RPMI 1640 medium containing 10% fetal bovine serum and 1% penicillin/streptomycin and incubated at 37°C with 5% CO2. After two hours, non-adherent cells were removed, leaving monocytes. These were cultured with 35 ng/mL GM-CSF and autologous serum (added every third day) for nine days, leading to differentiation into macrophages, characterized by larger size and protruding appendages, as observed in Supplementary Figure 1. The purity of the differentiated macrophages was assessed using flow cytometry. Cells were stained with fluorochrome-tagged antibodies specific for CD11b and CD14 (Suppl. Table 1). Cells with high expression of CD11b and low expression of CD14 (CD11b^high^ CD14^low^) were identified as differentiated macrophages, as shown in Supplementary Figure 2. To ensure the viability of the cells, a trypan blue exclusion assay was performed. The results indicated that more than 90% of the differentiated macrophages were viable. These viable, differentiated macrophages were subsequently used for further experimental procedures.

### Estimation of Advanced glycation end products (AGE) in serum

Estimating advanced glycation end products (AGEs) was conducted using the enzyme-linked immunosorbent assay (ELISA) (CSB-E09412h, Cusabio). Serum samples were collected from study participants and stored at -80°C until analysis. Standards and samples were prepared, with serial dilutions of the AGE standards creating a standard curve for quantification. Serum samples were diluted and added to microtiter wells pre-coated with AGE-specific antibodies. After a two-hour incubation at room temperature, wells were washed, and a biotinylated secondary antibody was added, followed by streptavidin-HRP conjugate. HRP substrate was then applied, producing a color change proportional to AGE levels. The reaction was stopped with sulfuric acid, and absorbance was measured at 450 nm. AGE concentrations were calculated using a standard curve generated from serially diluted AGE standards.

### Phagocytic activity of macrophages using FITC labeled BCG

The phagocytic activity of differentiated macrophages was assessed using a fluorescence-based method (11). BCG was labelled with FITC by incubating it with 5 mg/mL FITC for one hour at room temperature, followed by PBS washes to remove unbound dye. Macrophages were exposed to FITC-labelled BCG at an MOI of 10 for one hour at 37°C. To stop phagocytosis, cold PBS was added, and cells were washed thrice to remove free bacteria. Macrophage purity was confirmed by CD11b-PE-Cy7 and CD14-PerCP-Cy5.5 staining (Suppl. Table 1). To distinguish internalized from surface-bound bacteria, trypan blue was added to quench surface FITC fluorescence. Flow cytometry was used to measure the percentage of macrophages containing phagocytosed bacteria and their median fluorescence intensity (MFI).

### Surface expression of receptor for advanced glycation end products (RAGE) using flow cytometry

Surface expression of RAGE was estimated using rabbit anti-human RAGE antibody (Supp. Table 1). Briefly, 1x10^5^ macrophages were stained with the titrated volume of antibody for 30 minutes at 37^0^C. After incubation, cells were washed with FACS buffer (PBS + 0.5% BSA) and then stained with FITC-tagged Goat anti-rabbit antibody. After staining, cells were washed again and acquired on BD LSR fortessa X-20 and median fluorescence intensity (MFI) values and percentage positivity were recorded. Analysis was performed on FlowJo V10. Unstained controls were used as negative controls to control background autofluorescence. The experiment and analysis were performed according to the guidelines provided by Cossarizza et al used for flow cytometry and cell sorting in immunological studies(12).

### Separation of glycated and non glycated proteins and estimation of glycated calmodulin and glycated iNOS

After differentiation of monocytes to macrophages, the cells were lysed using Radioimmunoprecipitation Assay (RIPA) to obtain whole protein. The RIPA buffer contained components like Tris-HCl for stabilization, NaCl to prevent aggregation, NP-40, SDS, and Na-deoxycholate for protein extraction, and sodium azide to inhibit bacterial growth. About 1–2 million cells were processed with 200–300 µL RIPA buffer, vortexed, sonicated at 4°C, and centrifuged to collect the protein. Protein concentration was measured using the Bradford method. Glycated proteins, containing sugar groups, were separated using boronate affinity chromatography, which binds glycated proteins through their diol groups. Non-glycated proteins were collected in the flow-through, while glycated proteins were eluted with sorbitol.

Following separation of glycated and non-glycated proteins, In both glycated and non-glycated protein fractions, Calmodulin (CAM) and iNOS levels were measured using sandwich ELISA. For this, 100 µl of 0.8-1 µg/ml of total protein lysate from each fraction was used. The estimation was conducted using a commercially available ELISA kit (Supp. Table 1).

### Molecular Dynamics Simulations of CaM-iNOS-FMN complex

The glycosylated CaM-iNOS-FMN complex was generated using the crystal structure of Human iNOS Reductase and Calmodulin Complex (PDB ID:3hr4). The N-linked glycosylated lysines K21, K75, K77 and K94 (Figure 8b) in the CaM and the residues K516 and K531 in the iNOS-helical region were modeled manually by docking the sugar moiety N-acetyl glucosamine to an optimum covalent bond distance. The complex was refined, and stability of the protein was validated in the presence of the solvent. The MD simulation of the complex was carried out using Desmond (v-4.0) simulation package(13) in the presence of the TIP3P solvent model (14) using the OPLS 2005 force field parameters (15). Dimensions of simulation box (orthorhombic) extended by 10.0 Å in each direction from protein atoms. The system was neutralized by adding Na^+^ and Cl^-^ ions to balance the total charge for the system and bring it closer to the physiological environment (16). Simulations were performed with isothermal–isobaric NPT ensembles generated at 300K with the Nose-Hoover chain thermostat method by isotropic coupling at constant temperature with the Martyna-Tobias-Klein barostat method (17) with the help of RESPA integrator (18) and a time step of 2 fs (femtosecond). During the simulation, the geometric constraints were applied using the SHAKE algorithm (19). MD simulation was performed for the timescale of 1000 ns; the trajectories were recorded for every 200 ps, ∼1000 frames. The structural analysis was performed using Pymol 2.1.

### Statistical analysis

Statistical analyses were conducted using GraphPad Prism 6 (GraphPad Software Inc., San Diego, CA, USA). Categorical variables were presented as counts and percentages, while continuous variables were described as mean (SD) or median (interquartile range [IQR]) following the assessment of normality. Given non-normal distributions, non-parametric tests were employed throughout the study. Correlations between variables were assessed using Spearman correlation. The Mann-Whitney U and Kruskal-Wallis tests were used for comparisons between two groups and among three groups, respectively. A significance level of pl<l0.05 (two-tailed) was considered statistically significant for all analyses conducted.

### Data and resource availability statements

The datasets generated during and/or analyzed during the current study are available from the corresponding author upon reasonable request

## RESULTS

### Elevated circulating levels of advanced glycation end products (AGEs) and surface expression of receptor for AGE (RAGE) were found on macrophages under diabetic milieu

To investigate the role of AGEs and their receptor RAGE in TB patients under uncontrolled diabetic conditions, serum AGE levels and RAGE expression were analyzed. AGE levels were significantly higher in PTB+DM (57.70 ± 20.31 µg/mL) and DM (62.84 ± 26.83 µg/mL) patients compared to PTB (23.48 ± 14.05 µg/mL) and controls (25.31 ± 12.11 µg/mL) (p<0.001), with similar levels between PTB+DM and DM groups (Fig. 1a). RAGE expression followed a similar pattern, being higher in PTB+DM and DM groups compared to PTB (p<0.0007 and p<0.002) and controls (p<0.0001) (Fig. 1b), with comparable levels between PTB+DM and DM groups.

**Figure 1.**
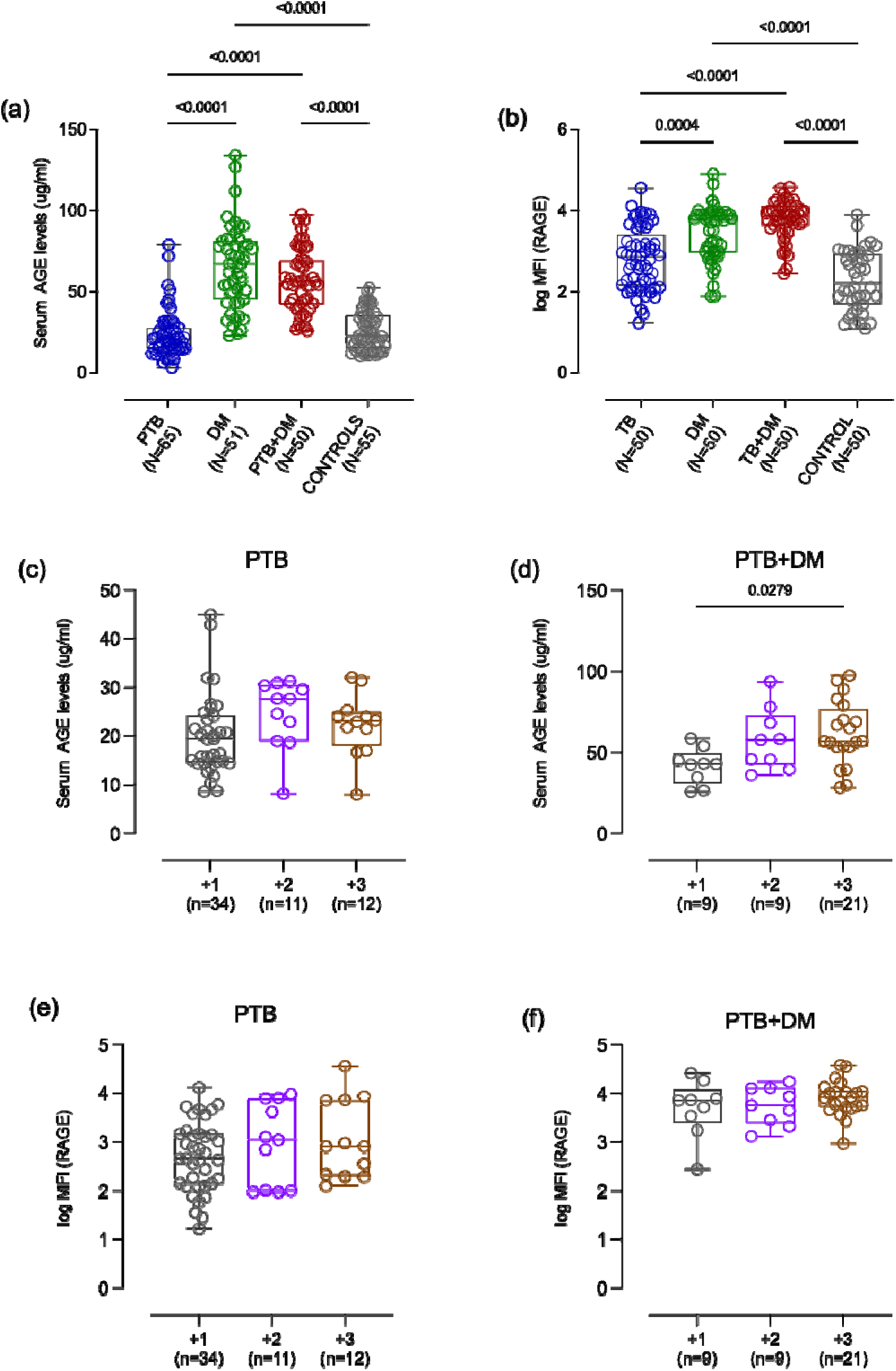
Serum advanced glycation end products (AGE) levels and receptor for advanced glycation end products (RAGE) expression on cultured macrophages in different study participants. NO was measured using ELISA. Surface expression of RAGE was measured by flow cytometry (a) shows serum AGE levels in PTB, DM, PTB+DM and controls. (b) shows the surface expression of RAGE in PTB, DM, PTB+DM and controls. (c) and (d) shows serum levels of AGE subdivided into different sputum grade PTB and PTB+DM patients. (e) and (f) show surface levels of RAGE on macrophages in PTB and PTB+DM subdivided into different sputum positivity. Data is represented as median with interquartile range and each point represents individual sample value. Box plot represents median with interquartile range. Kruskal-Wallis testing with post-hoc Dunn’s multiple comparison testing was performed. p values < 0.05 were considered to be statistically significant. One asterisk (*) indicates a p-value < 0.05; two asterisks (**) indicate a p-value < 0.01, three asterisks (***) indicate a p-value < 0.001 and four asterisks (****) indicate a p-value < 0.0001. PTB = Naïve active pulmonary TB; DM = Uncontrolled diabetic patients, PTB+DM = Uncontrolled diabetic patients with pulmonary TB, control = Healthy controls with no history of TB and DM.

AGE levels were significantly higher in 3+ sputum-positive PTB+DM patients than in 1+ (p<0.04), with no significant difference in PTB patients (Figure 1c and d). No statistical difference in RAGE expression was observed among different sputum grades (Fig. 1e and f).

### Advanced glycation end products (AGEs) positively correlated with RAGE and HbA1c in uncontrolled diabetic milieu with and without infection

AGEs binding to RAGE initiates various signalling cascade, generally pro inflammatory cascade upon binding to its ligands. In the present study, AGE is measured which is one of the ligands of RAGE. Therefore, we correlated levels of RAGE and AGE in PTB+DM and DM group. We found positive correlation between the receptor, RAGE and the ligand, AGEs (r=0.66 and 0.61 respectively) as shown in figure 2a and 2b. Both Advanced glycation end product (AGEs) and HbA1c (%) were higher in DM and PTB+DM group. Therefore, correlation analysis was done to find any correlation between both the parameters. As expected, a positive correlation was found between AGEs levels and HbA1c (%) in both the groups, PTB+DM (r=0.81) and DM (r=0.67) as shown in figure 2c and 2d. Similar findings were observed for surface expression of RAGE as shown in figure 2e and 2f.

**Figure 2.**
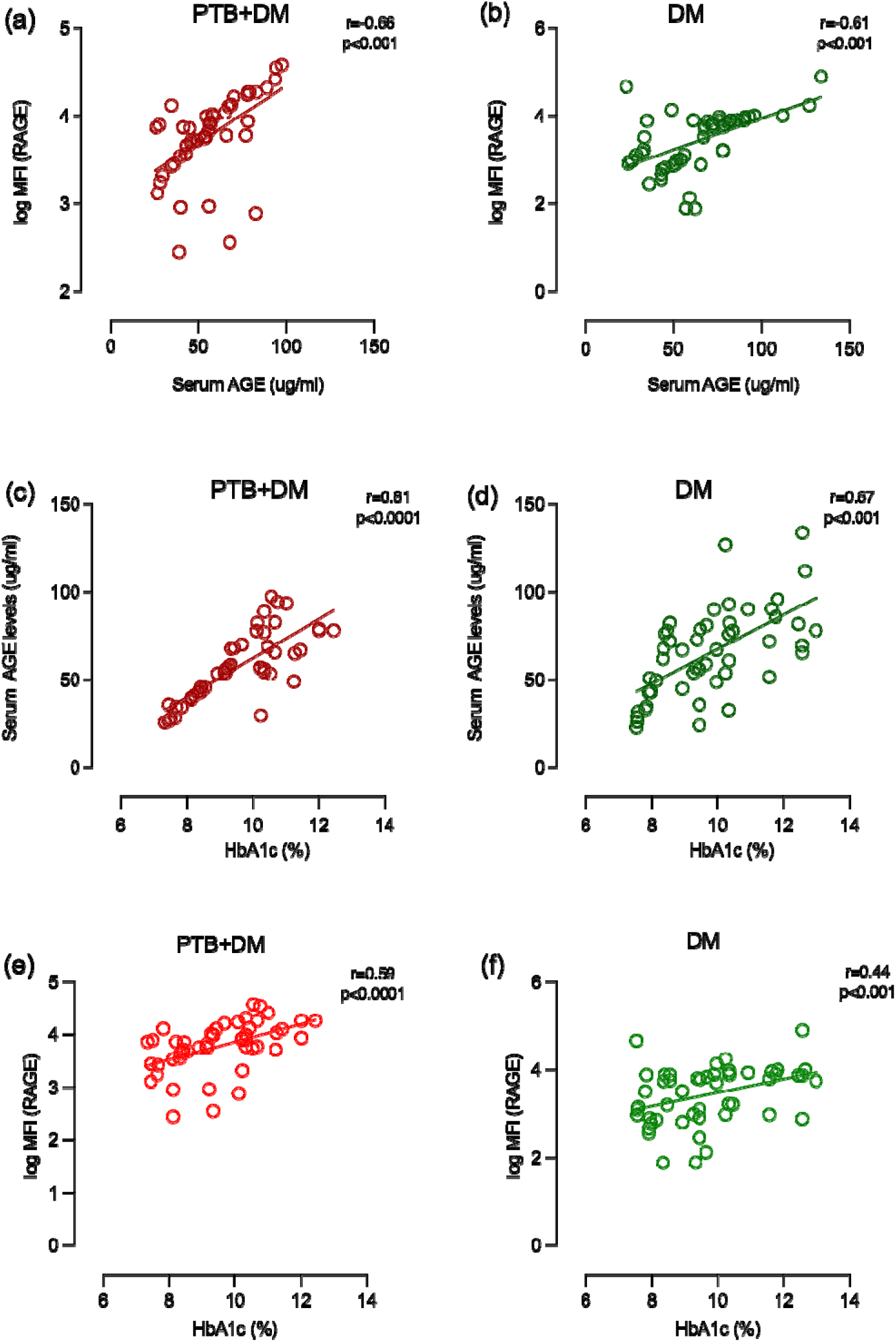
Correlation of serum AGE with surface expression of RAGE in (a) PTB+DM and (b) DM respectively. (c-f) shows the correlation of serum AGE and surface expression of RAGE with HbA1c levels in PTB+DM and DM respectively. Spearman correlation was used to correlate the variables. PTB+DM= Uncontrolled diabetic patients with pulmonary TB, DM = Uncontrolled diabetic patients. r = Spearman’s correlation coefficient. p shows the statistical significance. p-value less than 0.05 was considered significant

As the RAGE cascade leads to oxidative stress, we also wanted to check for an association between AGE and RAGE with ROS production in diseased group. We did not find a strong correlation between serum AGE levels and ROS production (figure 3a-c); however, RAGE was found to be positively correlated with ROS, suggesting higher oxidative stress as a result of higher RAGE expression (figure 3d-f).

**Figure 3.**
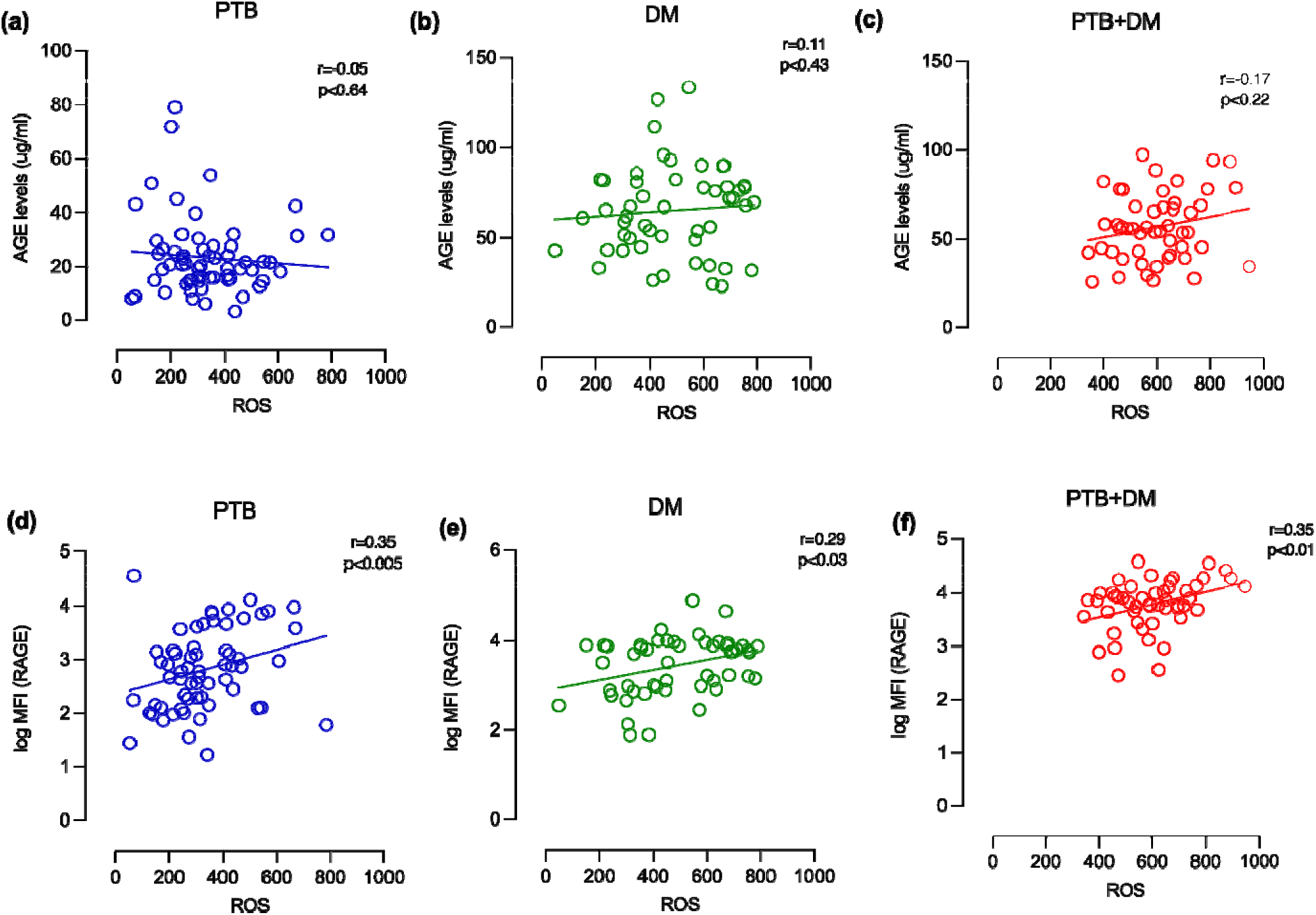
Correlation of serum AGE with ROS in (a) PTB and (b) DM and (c) PTB+DM respectively. (d-f) shows correlation of RAGE with ROS levels in PTB, DM and PTB+DM respectively. Spearman correlation was used to correlate the variables. PTB = Naïve active pulmonary TB; DM = Uncontrolled diabetic patients, PTB+DM= Uncontrolled diabetic patients with pulmonary TB. r = Spearman’s correlation coefficient. p shows the statistical significance. p-value less than 0.05 was considered significant

### Advanced glycation end products (AGEs) negatively correlated with phagocytosis capacity of macrophages

Advanced glycation end products have been shown to alter immune response. The presence of AGEs can impair the phagocytic activity of macrophages as shown by few previous studies. This impairment can reduce the ability to clear pathogens and cellular debris, contributing to chronic inflammation and tissue damage. To investigate the impact of AGEs on macrophage function, we analysed the correlation between AGE levels and macrophage phagocytosis across all study groups. Phagocytosis is a critical function of macrophages, essential for pathogen clearance and the initiation of appropriate immune responses. Previously, we reported lower phagocytic capacity of macrophages in diabetic milieu(11) . Here, we observed negative correlation between AGEs and phagocytosis index in all the patient groups as shown in figure 4.

**Figure 4.**
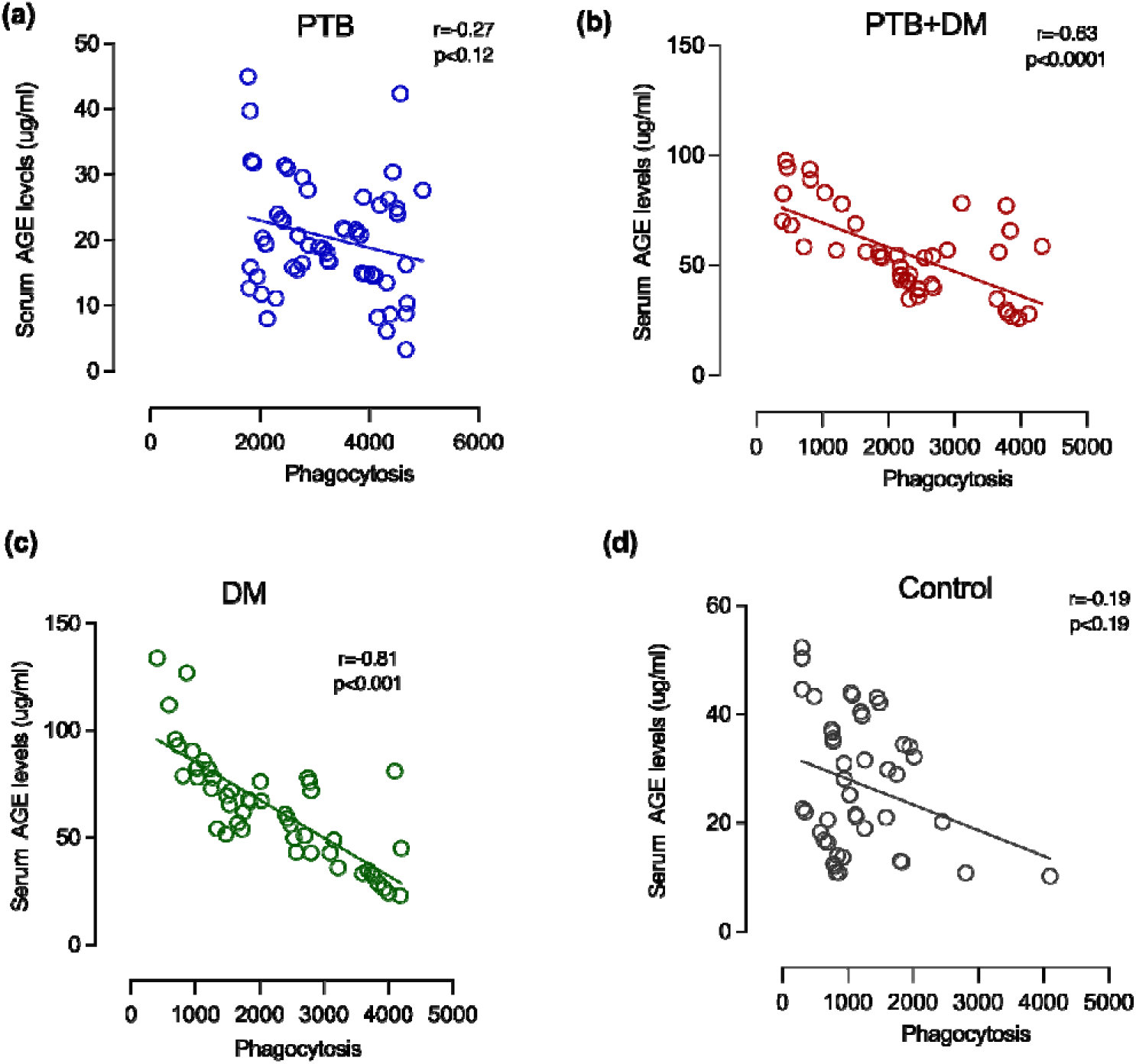
Correlation of serum AGE with phagocytosis. (a) shows correlation between AGE and HbA1c in PTB+DM and (b) DM, respectively. (c) shows correlation between RAGE and HbA1c in PTB+DM and (d) DM, respectively. Spearman correlation was used to correlate the variables. PTB = Naïve active pulmonary TB; DM = Uncontrolled diabetic patients, PTB+DM = Uncontrolled diabetic patients with pulmonary TB, control = Healthy controls with no history of TB and DM. r = Spearman’s correlation coefficient. p shows the statistical significance. p-value less than 0.05 was considered significant

### Higher glycated levels of calmodulin and iNOS in macrophages under uncontrolled diabetic milieu with and without infection

Along with AGEs, two glycated proteins namely, glycated calmodulin and glycated iNOS levels were estimated in macrophages of study participants since they are involved in NO production and their glycation under hyperglycemic condition may alter their function and hence impaired NO production. We observed a higher glycated calmodulin levels in PTB+DM and DM patients (4.49 ± 1.7 pg/mL and 4.04 ± 1.63 pg/mL respectively) as compared to PTB (2.55 ± 1.42 pg/mL) and healthy controls (2.44 ± 0.92 pg/mL) (p<0.01 and 0.001 respectively) as shown in figure 8. Glycated iNOS levels were also higher in PTB group (10.16 ± 4.10 pg/mL), DM group (7.99 ± 4.36 pg/mL) and PTB+DM group (9.79 ± 3.9 pg/mL) as compared to healthy controls (4.76 ± 2.05 pg/mL) with p<0.001 as shown in figure 5. However, no difference was observed across PTB, DM and PTB+DM group.

**Figure 5.**
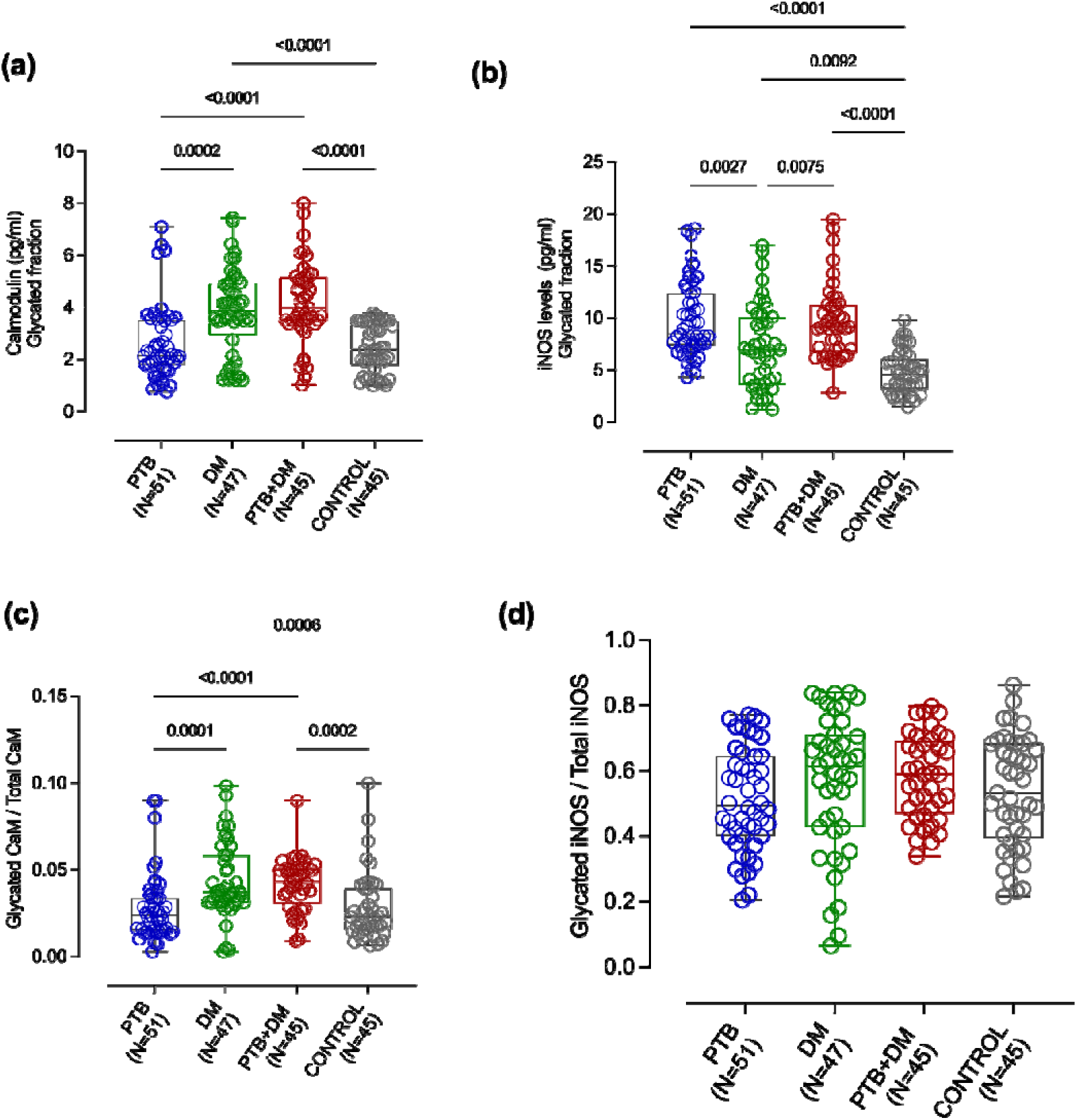
Glycated calmodulin (CAM) and glycated iNOS levels in different study participants. (a) represents total glycated CAM levels in PTB, DM, PTB+DM and controls. (b) represents total glycated CAM levels in PTB, DM, PTB+DM and control, (c) and (d) represents fraction of glycated CAM and iNOS respectively. Data is represented as median with interquartile range and each point represents individual sample value. Box plot represents median with interquartile range. Kruskal-Wallis testing with post-hoc Dunn’s multiple comparison testing was performed. p values < 0.05 were considered to be statistically significant. One asterisk (*) indicates a p-value < 0.05; two asterisks (**) indicate a p-value < 0.01, three asterisks (***) indicate a p-value < 0.001 and four asterisks (****) indicate a p-value < 0.0001. PTB = Naïve active pulmonary TB; DM = Uncontrolled diabetic patients, PTB+DM = Uncontrolled diabetic patients with pulmonary TB, control = Healthy controls with no history of TB and DM.

### Glycated calmodulin negatively correlated with serum NO levels

Since glycated calmodulin levels were higher in diabetic milieu and can affect NO production via iNOS, we assessed the correlation between glycated calmodulin and NO in the study groups. A negative correlation was found between glycated calmodulin and NO in PTB+DM and DM group (r=-0.40 and -0.36 respectively) shown in figure 6a-6b). A weak correlation was found in PTB group as well (r=-0.25) while no correlation was found in healthy control group (figure 6c-6d). Unlike glycated CAM, glycated iNOS was not found to be correlated with serum NO levels except in diabetic individuals where the NO levels were negatively correlated with glycated iNOS levels (figure 6e-6h).

**Figure 6.**
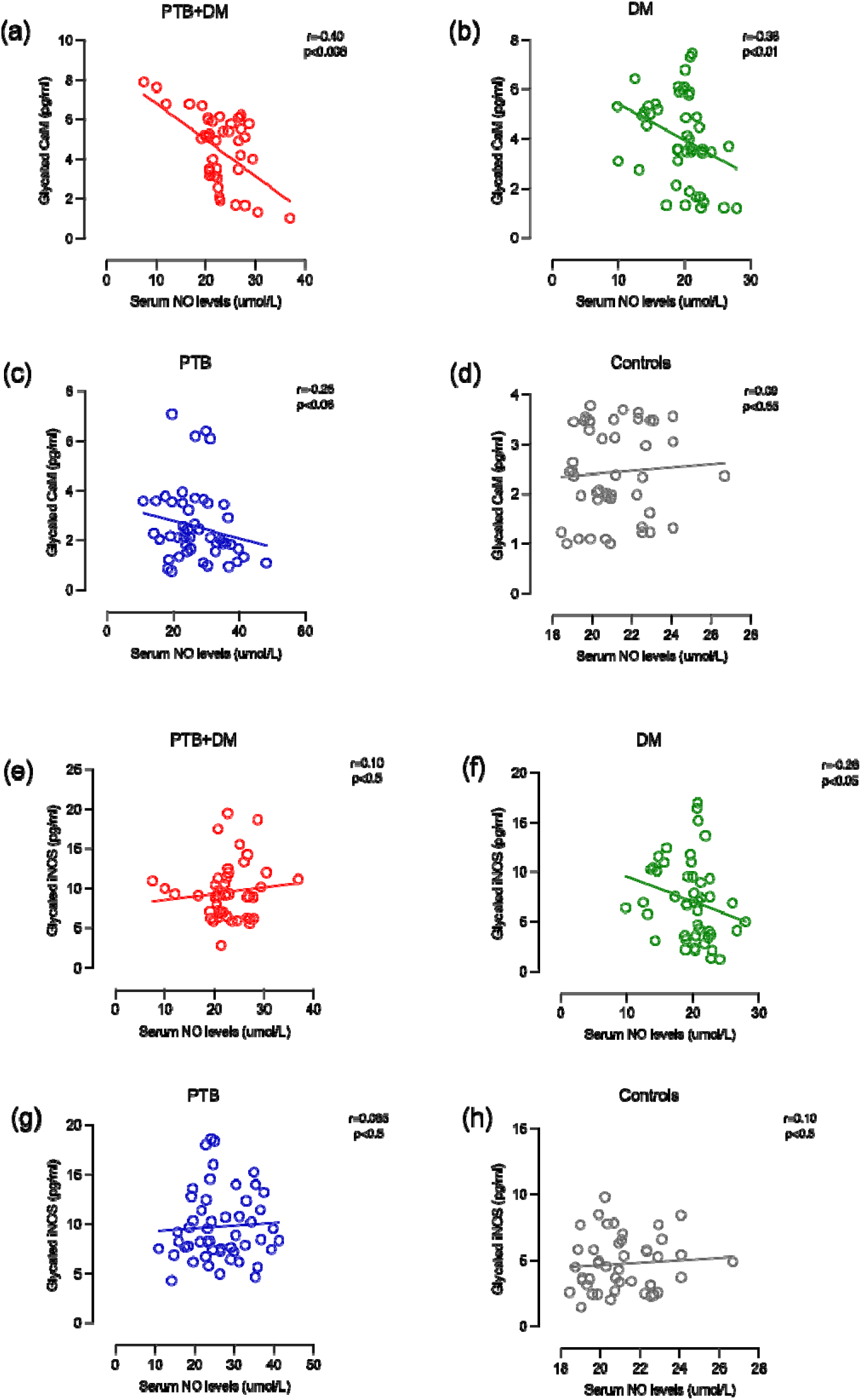
Correlation of glycated CAM with Serum NO levels. (a-d) shows correlation between glycated CAM with Serum NO levels in PTB, DM, PTB+DM and controls respectively. (e-h) shows correlation between glycated iNOS with Serum NO levels in PTB, DM, PTB+DM and controls respectively. Spearman correlation was used to correlate the variables. PTB = Naïve active pulmonary TB; DM = Uncontrolled diabetic patients, PTB+DM = Uncontrolled diabetic patients with pulmonary TB, control = Healthy controls with no history of TB and DM. r = Spearman’s correlation coefficient. p shows the statistical significance. p-value less than 0.05 was considered significant

We also looked for any correlation of glycated calmodulin and iNOS with HbA1c. We found no significant correlation between glycated calmodulin and iNOS with plasma HbA1c levels as shown in figure 7.

**Figure 7.**
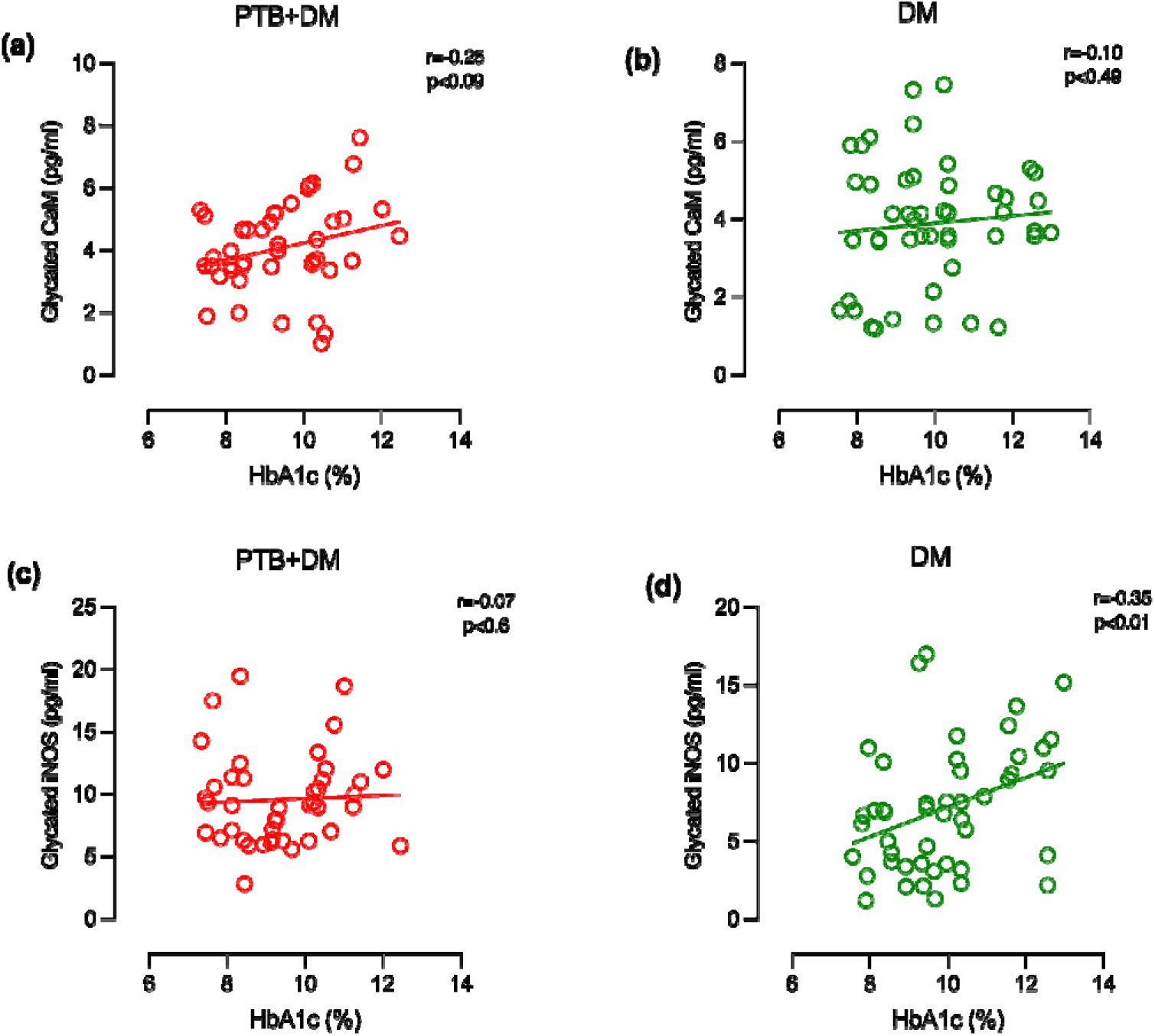
Correlation of glycated CAM and glycated iNOS with HbA1c levels. (a) shows correlation between glycated CAM with HbA1c levels in PTB+DM and (b) DM, respectively. (c) shows correlation between glycated iNOS with HbA1c levels in PTB+DM and (d) DM, respectively. Spearman correlation was used to correlate the variables. PTB+DM= Uncontrolled diabetic patients with pulmonary TB, DM = Uncontrolled diabetic patients. r = Spearman’s correlation coefficient. p shows the statistical significance. p-value less than 0.05 was considered significant

### Glycosylation of Calmodulin leads to reduced interaction between calmodulin and iNOS

The iNOS is an active homodimer consisting of the terminal oxidase domain and the C terminal reductase domain with sub-domains for the FAD/NADPH-binding and the FMN-binding (Figure 8a)(20). A 30-residue alpha-helical linker region called the CaM-recognition site connects the FMN subdomain to the oxidase domain (Figure 8a) (21). For catalysis and NOS production, the electron from the reductase domain is transferred to the heme molecule in the oxidase domain, which requires CaM to bind the linker region (22). The binding releases the FMN subdomain from the input to the output state to release electrons. Hence, the NO production is regulated by interdomain interaction between iNOS and calmodulin. Based on our data, we proposed that the glycosylated CaM and iNOS would have a lower affinity towards each other. To understand the proposed mechanism, we performed 1µs MD simulations of the iNOS-FMN domain-CaM complex (PDB ID: 3hr4_A) with glycosylated lysine (K21, K75, K77 and K94) (Figure 8b) in the N-and C-lobes of CaM and glycosylated lysine (K516 and K531) in the 30-residue alpha-helical linker region of iNOS (Figure 8b). The MD trajectory showed that the complex is unstable, losing significant interactions between the linker helix and CaM domains. The non-glycosylated complex is majorly stabilized by hydrophobic, salt-bridge, and H-bonded interactions, as shown in Figure 1a (23). The 1µs MD trajectories of the complex revealed a complete loss of all salt-bridge and H-bonded interactions between iNOS-FMN and CaM domains. The superposition of the complex before and after 1µs MD run showed significant movement of the iNOS-FMN domain from CaM interface (Figure 8c). Similarly, the N-and-C-lobe of CaM significantly lost van der Waals contact with the iNOS-linker region, as shown in Figure 8d. Although we did not observe complete dissociation of the complex, we could observe that if we run MD for a more extended period, the complex might dissociate. Hence, we can propose that glycosylation interferes with the calmodulin binding to iNOS and will impact the enzyme’s production of NO.

**Figure 8:**
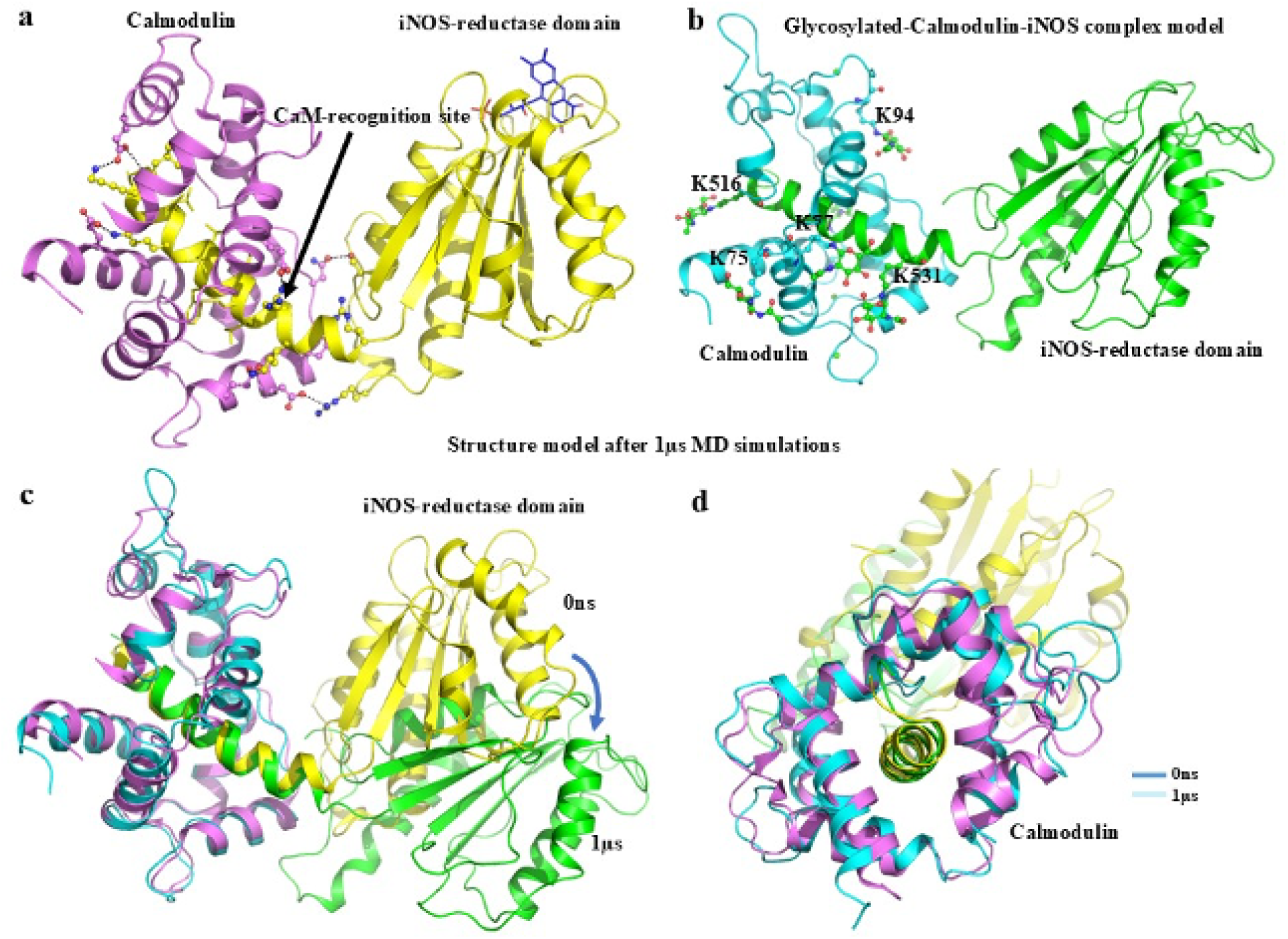
(a) iNOS-reductase domain in complex with CaM (PDB ID: 3hr4) showing the residues forming salt-bridge and H-bonded interactions in ball-and-stick. (b) the Model complex of glycosylated iNOS-FMN domain and CaM. The residues linked with N-acetyl glucosamine sugar moiety (K21, K75, K77, and K94) in CaM and (K516, K531) (iNOS linker region) are shown in ball-and-stick. (c,d). Superposition of iNOS-FMN domain-CaM complex before (0ns, blue) and after 1µs (cyan) MD simulations showing the movement of iNOS-FMN domain in the glycosylated complex. The dissociation of the CaM domain from the iNOS helical region is observed due to glycosylated lysine residues (K516 and K531).

## DISCUSSION

During prolonged hyperglycemia in diabetes mellitus, glucose undergoes non-enzymatic glycation with plasma proteins, leading to the formation of advanced glycation end products (AGEs) (4,24). These AGEs play a critical role in the pathogenesis of diabetic complications by binding to the receptor for AGEs (RAGE) (6,24,25). The AGE-RAGE interaction initiates signaling cascades that trigger inflammation, the release of pro-inflammatory molecules, and the generation of free radicals (6,26). This mechanism significantly modulates immune responses, including those involving macrophages, which are crucial for defending against infections like tuberculosis (TB). We previously demonstrated a dynamic link between diabetes mellitus and tuberculosis, highlighting impaired macrophage effector functions. (11). In the present study, we investigated AGE and RAGE levels in patients with pulmonary TB who also have diabetes (PTB+DM), compared to those with only pulmonary TB (PTB), only diabetes (DM), and healthy controls. We found that both AGEs and RAGE levels were significantly elevated in diabetic individuals, regardless of TB status, indicative of prolonged exposure to hyperglycemia. Our findings align with a 2019 study reporting elevated soluble RAGE (sRAGE) and RAGE ligands in PTB-DM and DM compared to PTB and household contacts (7). Another study showed no difference in CML, a type of advanced glycation end product, but they found higher levels of soluble RAGE in TB patients showing their association with disease pathology (27). These ligand-receptor levels were also positively correlated in the diabetic milieu. The binding of AGEs to RAGE creates a positive feedback loop, where the activation of RAGE signaling pathways further enhances the production and accumulation of AGEs (8,28). Elevated AGEs in diabetic conditions indicate irreversible protein adduct formation through non-enzymatic glycation, likely impairing macrophage function via RAGE binding. Our findings of higher AGE levels in patients with increased sputum positivity suggest AGEs may affect disease severity. While RAGE levels did not differ significantly among sputum-positive patients, the higher trend in PTB cases suggests TB infection may influence RAGE expression, impacting immune responses. AGE-RAGE interactions on macrophages likely trigger pro-inflammatory cytokines and oxidative stress (29,30). This could explain our findings of a positive correlation between RAGE and reactive oxygen species (ROS) production in individuals with diabetes and TB infection. Previous research in experimental pneumonia has shown that RAGE deficiency correlates with improved survival and reduced bacterial dissemination, where mice lacking RAGE had a better survival rate together with a lower pulmonary bacterial load and decreased dissemination of S. pneumoniae to blood and spleen compared to wildtype mice (31) , highlighting the potential detrimental role of RAGE in infectious diseases. Interestingly, we found significant correlation of HbA1c with both AGE and RAGE, suggesting that higher blood glucose levels lead to an increase in these markers. In light of our findings, we hypothesized that long-term uncontrolled diabetes leads to a higher RAGE level, enough to cause detrimental effects. Our study found an inverse correlation between AGE levels and macrophage phagocytic capacity, aligning with prior evidence of AGE-induced suppression of macrophage function. AGEs impair phagocytosis and cytokine production, weakening host defense mechanisms (32). Another recent study also showed that glycation contributes to changes in macrophage activity and cytokine expression and, therefore, could support the understanding of disturbed wound healing during aging and diabetes(33). Chronic exposure to AGEs may, therefore, compromise the ability of macrophages to mount effective immune responses against infections, contributing to reduced phagocytic activity and overall immune dysregulation.

Macrophage phagocytic activity is essential for breaking down and removing engulfed bacteria, relying on reactive oxygen species (ROS) and reactive nitrogen species (RNS). RNS synthesis requires the enzyme inducible nitric oxide synthase (iNOS) (34,35) . iNOS typically exists as a monomer and is activated by dimerization, facilitated by calmodulin (CAM)(36).

This triggers nitric oxide (NO) production, crucial for bacterial killing. In chronic hyperglycemia, such as in diabetes, there is an increased glycation of proteins within macrophages and in the serum. Glycation is a non-enzymatic process where glucose molecules attaches to proteins, altering their structure and function (37) . The CAM protein, which has free lysine and arginine chains, is highly prone to glycation in hyperglycemia. iNOS can also become glycated, possibly impairing its role in producing NO, critical for killing bacteria. Therefore, we investigated glycation in CAM and iNOS in macrophages. Comparable glycated CAM levels were found in pulmonary TB (PTB) patients and healthy controls. However, PTB+DM and DM patients had higher glycated CAM levels, likely due to prolonged hyperglycemia. While glycated iNOS levels did not differ significantly between groups, they were elevated in patient groups compared to controls, likely due to infection and inflammation. Our findings suggest that increased glycation of CAM under hyperglycemic conditions may impair its ability to activate iNOS effectively, leading to reduced NO production. Therefore, our next step was to see the effect of these glycation on NO production. A negative correlation between glycated CAM and NO levels was observed, supporting the hypothesis that glycated CAM reduces NO production. Glycated iNOS showed no significant correlation with NO, suggesting CAM plays a larger role in this effect. Reduced NO may weaken macrophages’ ability to kill bacteria, increasing infection risks in diabetic patients.

Considering the significant alterations in glycated calmodulin levels in a diabetic environment, we evaluated its correlation with iNOS and HbA1c levels. Although a positive trend was observed, it lacked statistical significance, potentially due to biological variability or limited sample size. Future studies with refined designs are needed to establish HbA1c thresholds at which glycated calmodulin impacts iNOS function and NO production critical for bacterial killing.

High Glycated CAM in diabetic milieu observed earlier may lead to structural changes, reducing its binding efficiency to iNOS and thereby hampering NO production. To address this, molecular dynamic studies were carried out to understand the effect of glycosylation on interaction of iNOS and calmodulin. We performed MD simulations of the iNOS-FMN domain-CaM complex with glycosylated lysine (Residue K21, K75, K77 and K94) in the N- and C-lobes of CaM and glycosylated lysine (Residue K516 and K531) in the 30-residue alpha-helical linker region of iNOS. This analysis showed the complex is unstable, losing significant interactions between the linker helix and CaM domains. Additionally the non-glycosylated complex is majorly stabilized by hydrophobic, salt-bridge, and H-bonded interactions further validating our hypothesis that glcosylation of iNOS and calmodulin disrupt NO production.

This study offers insights into mechanisms behind increased tuberculosis susceptibility in type 2 diabetes but has limitations. It included only uncontrolled diabetics, excluding pre-diabetic and controlled cases, limiting understanding across hyperglycemia levels. Additionally, other soluble RAGE ligands, which may impact disease outcomes, were not evaluated.

In summary, our study emphasizes the role of advanced glycation end products (AGEs) and their receptor (RAGE) in immune dysregulation in diabetic patients, especially those with pulmonary tuberculosis (PTB). Chronic hyperglycemia leads to protein glycation, forming AGEs that activate pro-inflammatory pathways and oxidative stress through RAGE, impairing macrophage function (low phagocytic activity and higher oxidative stress) and worsening disease severity. High glycated CAM and iNOS levels in uncontrolled hyperglycemia interfere with nitric oxide production, affecting infection control. These findings highlight the need for therapies targeting AGE accumulation or AGE-RAGE signaling to restore immune responses and reduce infection risk in diabetic patients. Future research should explore AGE-RAGE mechanisms and HbA1c thresholds to guide new treatments for managing diabetes and related infections like TB.

## Acknowledgements

We thank the DOTS center, AIIMS, and Safdarjung Hospital and all the study subjects for participation in the study. We thank the Science and Engineering Research Board (SERB), India”, SERB-DST for providing the research grant for the study (EEQ/2017/000165).

## Author Contributions

Archana Singh and Sudhasini Panda conceptualized and designed the study. Sudhasini Panda drafted the manuscript. Sudhasini Panda and Diravya M Seelan carried out recruitment of patients under the guidance of Archana Singh, Anant Mohan, Naval K Vikram, and Neeraj Kumar Gupta. Sudhasini Panda and Diravya M Seelan carried out sample collection, standardization, and execution of experimental work along with data acquisition and interpretation of data under the guidance of Archana Singh and Kalpana Luthra. Alisha Arora, Gunjan Dagar executed some of the experimental work along with data acquisition. Alankrita Singh and Abdul S Ethayathulla carried out molecular Dynamic studies. Archana Singh and Mayank Singh critically reviewed and contributed to the final version of the manuscript. Archana Singh gave the final approval of manuscript submission and supervised the entire project.

## Guarantors

Archana Singh is the guarantor of this work and, as such, had full access to all the data in the study and takes responsibility for the integrity of the data and the accuracy of the data analysis.

