## Supplementary table 1 for "Implications of diabetes mellitus on pathophysiology of Tuberculosis: Role of AGE-RAGE Axis and Glycated Proteins"

| Reagents | | |  |
| --- | --- | --- | --- |
| Flow antibodies | | |  |
| Marker | **Catalog number** | **Company** | **RRID** |
| CD14/ PerCP cy5.5 | 562692 | BD Biosciences | AB_2737726 |
| CD11b/ PE Cy7 | 557743 | BD Biosciences | AB_396849 |
| RAGE | OAAB02276 | Aviva Systems Biology | AB_3668793 |
| FITC tagged Goat anti-rabbit antibody | A-11011 | Thermo Scientific | AB_143157 |
| ELISA kits | | |  |
| Protein | **Catalog number** | **Company** |  |
| Human Advanced glycation end products | CSB-E09412h | Cusabio |  |
| Human Calmodulin | E-EL-H0625 | Elabsciences |  |
| Human iNOS | E-EL-H0753 | Elabsciences |  |
