## Supplementary material for "Implications of diabetes mellitus on pathophysiology of Tuberculosis: Role of AGE-RAGE Axis and Glycated Proteins": Supplememntary figure 1

**Supplementary Data**

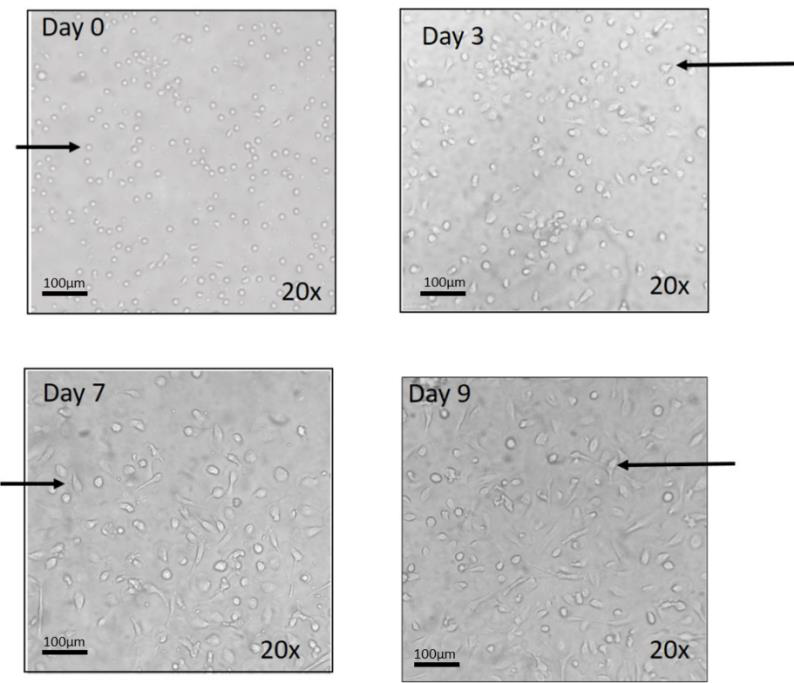

**Supplementary figure 1** – Morphological changes of differentiating macrophages. Monocytes were differentiated into macrophages using GM-CSF. Morphological changes of differentiated macrophages were visualized by phase contrast inverted microscope under 20X magnification at different time interval of (a) day 0, (b) day 3, (c) day 7 and (d) day 9. Differentiated macrophages were increased in size and had protruding appendages. Scale- 100μm
