## Supplementary figure 2 for "Implications of diabetes mellitus on pathophysiology of Tuberculosis: Role of AGE-RAGE Axis and Glycated Proteins"

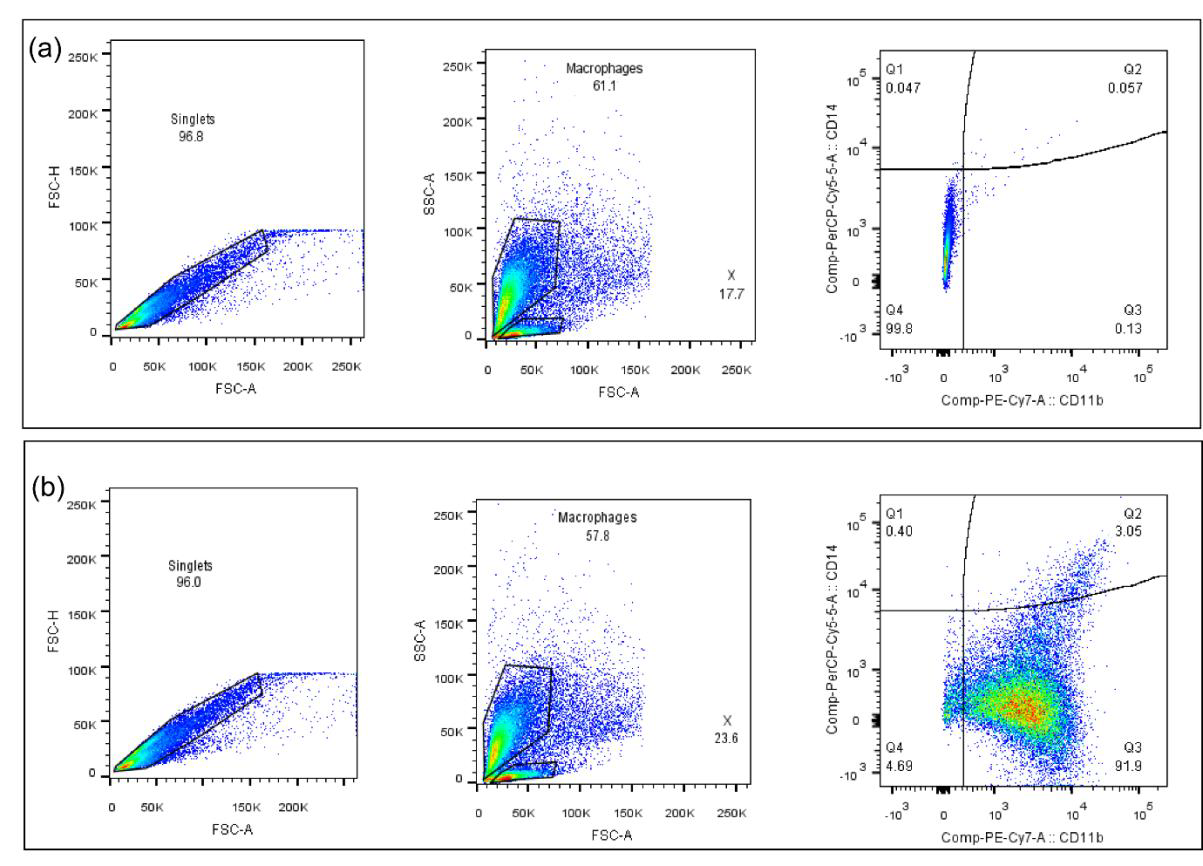

**Supplementary figure 2 –** Pseudocolor plot showing sequential gating strategy used to identify differentiated macrophages (a) unstained cells. (b) Cells stained with CD14 and CD11b. Ungated events were initially plotted as FSC-A × FSC-H for selection of single cells and exclusion of cell aggregates located far from the main diagonal. morphologically similar cells were represented in SSC-A × FSC-A plots. Then these cells were gated for CD11b and CD14. Differentiated macrophages were shown as CD11bhigh CD14low.
